# CD44 regulates blood-brain barrier integrity in response to fluid shear stress

**DOI:** 10.1101/2020.01.28.924043

**Authors:** Brandon J. DeOre, Paul P. Partyka, Fan Fan, Peter A. Galie

## Abstract

Fluid shear stress is an important mediator of vascular permeability, yet the molecular mechanisms underlying the response of the blood-brain barrier to shear have yet to be studied in cerebral vasculature despite its importance for brain homeostasis. The goal of this study is to probe components of shear mechanotransduction within the blood-brain barrier to gain a better understanding of pathologies associated with changes in cerebral blood flow including ischemic stroke. Interrogating the effects of shear stress in vivo is complicated by the complexity of factors in the brain parenchyma and the difficulty associated with modulating blood flow regimes. Recent advances in the ability to mimic the in vivo microenvironment using three-dimensional in vitro models provide a controlled setting to study the response of the blood-brain barrier to shear stress. The in vitro model used in this study is compatible with real-time measurement of barrier function using transendothelial electrical resistance as well as immunocytochemistry and dextran permeability assays. These experiments reveal that there is a threshold level of shear stress required for barrier formation and that the composition of the extracellular matrix, specifically the presence of hyaluronan, dictates the flow response. Gene editing to modulate the expression of CD44, a receptor for hyaluronan that previous studies have identified to be mechanosensitive, demonstrates that the receptor is required for the endothelial response to shear stress. Manipulation of small GTPase activity reveals CD44 activates Rac1 while inhibiting RhoA activation. Additionally, adducin-γ localizes to tight junctions in response to shear stress and RhoA inhibition and is required to maintain the barrier. This study identifies specific components of the mechanosensing complex associated with the blood-brain barrier response to fluid shear stress, and therefore illuminates potential targets for barrier manipulation in vivo.

## Introduction

Recent studies have established that the integrity of the blood-brain barrier (BBB) is regulated by mechanical stress exerted by blood flow; both fluid shear stress^1^ and cyclic strain^2^ increase the barrier function of in vitro models of cerebral vasculature. Therefore, an understanding of how the BBB responds to mechanical stress has relevance to neurolopathologies associated with changes to the vasculature and blood flow. One such example is ischemic stroke: an occlusion reduces blood flow prior to reperfusion, and subsequently a no-reflow period occurs as cerebral blood flow is attenuated while vascular permeability simultaneously increases ^3–5^. Given that previous studies have demonstrated that shear stress stabilizes the barrier, it is likely that the reduction in blood flow causes or at least contributes to BBB breakdown during the no-reflow period. Yet despite its importance in disease and overall brain homeostasis, the mechanisms underlying the BBB response to fluid shear stress have not yet been investigated in detail.

Several mechanosensing mechanisms of fluid shear stress have been identified in systemic vasculature^6–9^, including pathways involving the glycocalyx ^10^, the PECAM-VEGFR2-VE-cadherin complex ^11^, and Notch signaling ^12^. Yet the differences in structure and function of vasculature and surrounding extracellular matrix in the central nervous system (CNS) insinuate the potential for unique mechanotransduction pathways in the BBB. Endothelial cells in cerebral vasculature form tight junctions, which are complexes of claudins and occludens stabilized by scaffolding proteins including zonula occludin that give rise to the BBB and prevent the transport of blood-borne solutes across the vessel wall into the brain parenchyma. Moreover, the extracellular matrix (ECM) of the CNS is drastically different than other organ systems. The ECM features high concentrations of proteoglycans and glycosaminoglycans, specifically hyaluronan (HA)^13^. Given these differences and the wide variety of mechanotransduction pathways that have been identified in system vasculature^14^, the present study interrogates a novel mechanism relevant to the BBB.

One common component in several shear-mediated signaling pathways involves the activation of small GTPases^8, 15^. One small GTPase in particular, RhoA, is also implicated as a regulator of BBB integrity^16–19^. Previous studies have demonstrated that increased RhoA activity leads to a disassembly of the complexes within cell-cell junctions and subsequent barrier disruption^8, 20, 21^. Another small GTPase, Rac1, has been associated with stabilization of the BBB^22, 23^, and recent studies have also identified Rac1 activation as a downstream response to fluid shear stress^12^. Therefore, this study investigates the effect of shear stress on the activity of both RhoA and Rac1 to gain insight into their role in shear-mediated barrier integrity.

The results described here were produced by a previously described three-dimensional model of the BBB^2^. Despite several differences compared to in vivo vasculature, including the lack of a tortuous and branched morphology, absence of immune cells including microglia, and use of culture medium for perfusion instead of blood, the in vitro model provides a controlled setting to study the response to shear stress. Moreover, cells can be genetically or transcriptionally altered prior to incorporation into the device to determine the effects of specific components of the signaling pathway. The device is compatible with dextran-FITC permeability assays as well as immunocytochemistry. A TEER device has also been developed for this microfluidic system, allowing non-invasive measurements of barrier integrity. Overall, the device provides a robust platform to study BBB mechanotransduction.

## Methods

### Microfabrication

Polydimethylsiloxane (PDMS) microfluidic devices were manufactured as described previously^2^. Briefly, positive master molds of the devices manufactured using stereolithography (Protolabs) were used to cast negative and positive PDMS master molds which were then used to fabricate the microfluidic device. For monolayer flow experiments, 5-g of PDMS was added to a p100 culture dish and leveled prior to curing. The hydrogel reservoir was etched with 5M sulfuric acid for 90 minutes, thoroughly washed with Millipore water, and subsequently coated with 20 µg/mL collagen for 60 minutes. For monolayer plates, a 40-mm glass cover slip was used to constrain the fluids on the dish. All steps were performed at room temperature. Devices were sterilized under shortwave length ultraviolet light prior to cell seeding.

### Cell culture

All cell experiments were performed with p21-24 HCMEC/d3 (gifted from Dr. Robert Nagele’s lab at the Rowan School of Osteopathic Medicine) 3-5 days after thawing and feeding with modified EGM-2 on 1% gelatin coated tissue culture plates^24^. Normal human astrocytes (NHA)(Lonza) were thawed at P5 and cultured for 5-10 days prior to cell seeding. Cell cultures were maintained at 37°C with 5% CO^2^ and 95% relative humidity.

### In vitro blood-brain barrier model

Three-dimensional models of the blood-brain barrier (3D BBB) were fabricated as described previously^2^. Briefly, a hydrogel composed of 5 mg/mL type I collagen, 1 mg/mL HA, 1 mg/mL Matrigel^25^ were used to fabricate the scaffold. When incorporated into the hydrogel, NHA were seeded at 1 million cells per mL. The gel was injected into the hydrogel reservoir of the device and 180-µm needles coated in 0.1% BSA were inserted prior to polymerization of the hydrogel. The needles were removed after the gel was polymerized leaving two voids in the hydrogel in which HCMEC/d3 were injected into one channel at a density of 10 million per mL (15 µL per channel). Channels were incubated for 10 minutes to ensure cell attachment then injected with cells again and inverted for 10 minutes to coat the opposite side. Following cell seeding, channels were either exposed to flow using a linear syringe pump (Kent Scientific) or incubated in static conditions in a 6-well plate. For GTPase activity assays, a larger diameter (1-mm) model was used, which was characterized in previous work^26^. For the monolayer experiments, the same hydrogel formulation was polymerized on the treated PDMS coated plates under a sterile 40-mm glass coverslip to create a uniform circular hydrogel. Following removal of the cover slip, HCMEC/d3 were seeded on the gel at a density of 4k/cm^2^ and allowed to adhere for 30 minutes prior to addition of EGM-2. Monolayers were incubated for 4 days in static culture to ensure confluency, then exposed to fluid shear stress. Fluid shear stress was applied using a 40-mm 1-degree cone plate on a rheometer (Waters) for 24-hrs on a Peltier plate set to 37°C. The media was supplemented with HEPES buffer to a final concentration of 10 mM to maintain pH and sterile Millipore was added to the plate during exposure of flow to counteract evaporative loss^27^.

### Immunocytochemistry

Following exposure to experimental conditions, vessels were fixed in 4% paraformaldehyde (Alfa Aesar) for 30 min at room temperature. Following fixation, the top layer of the device was removed with a razor blade and then the hydrogel was removed from the device and placed in 0.1% Triton X-100 to permeabilize the cell membrane. Gels were blocked in 5% normal donkey serum or 3% BSA for 30 minutes at room temperature followed by incubating overnight with primary antibodies for either HCAM (CD44)(Santa Cruz), glial fibrillary acidic protein (GFAP), adducin-γ (ADD3)(ABcam), or zonula occludin-1 (ZO-1)(CST). Following primary incubation, gels were washed three times with PBS for 5 minutes then incubated with the appropriate secondary antibody conjugated to Alexa 555 (CST), Alexa 488(Santa Cruz), or Dylight 650 (Thermo Scientific). Gels were counter-stained with DAPI to label nuclei and FITC-phalloidin for actin. All gels were imaged using a Nikon A-1 confocal scanning microscope.

### Permeability testing

After exposure to experimental conditions, channels were transferred to the stage of an inverted epifluorescent microscope enclosed by an environmental chamber set to 37C, 5% CO2, and 95% RH. The channels were perfused with 4-kDa dextran-FITC at a flow rate of 5 µL/min using a syringe pump for 10 minutes, while submerged within culture medium to ensure cell viability. This flow rate was selected to assure fully developed flow throughout the channel and to maintain consistency with previous work^2^. Images were taken at 30s intervals for 10 minutes, and the diffusion coefficients were established using the following equation from previous work^28^.

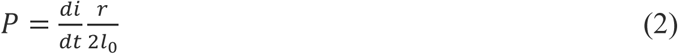

Sample numbers of at least n = 3 were used to determine the mean and standard deviation of diffusion coefficients for each condition.

### Impedance spectroscopy and transendothelial electrical resistance (TEER) measurements

Impedance measurements were obtained a stingray DS1M12 USB oscilloscope adapter (USB Instruments) that measured current across a reference resistor for a range of frequencies. Impendence, defined as Z=V/I, was measured at 15-Hz, where capacitance of the electrodes dominates and 15.6-kHz, where resistance of the culture media dominates^29^; the difference between these two values yielded the TEER measurement. These values were normalized using the impedance of an acellular hydrogel within the microfluidic device.

### CD44 knockdown and upregulation

Commercially available CRISPR plasmids (Santa Cruz) were used to alter the expression of CD44 within the HCMEC/D3 cells prior to introduction into the 3D BBB model. To knockout CD44 in the cells, two plasmids encoding a D10A mutated Cas9 nuclease and a CD44-specific 20 nucleotide guide RNA were transfected into cells (sc-400209-NIC). The paired guide RNA sequences were offset by approximately 20 basepairs to facilitate Cas9-mediated double nicking of genomic DNA. For scrambled controls, a single plasmid encoding a non-specific guide RNA sequence was transfected into cells (sc-418922). For the upregulation condition, lentiviral activation particles containing a SAM complex were delivered to cells to activate transcription of CD44 (sc-400209-LAC). Protocols for these assays were adapted from instructions provided by Santa Cruz. Briefly, cells were plated in a six well plate and grown to 60-70% confluency. For HCAM activation, the cells were incubated with complete media containing 10 µg/mL polybrene prior to adding lentiviral particles at an MOI of 2. After 2 days, stable colonies were selected by incubation with 5 µg/mL puromycin in EGM-2. Cells were transfected with KO and control plasmids resuspended in nuclease free water at 0.1 µg/µL. Both solutions were incubated for 5 minutes at room temperature. Following incubation, solutions were mixed, vortexed, and incubated for 20 minutes at room temperature. 300 µL of the plasmid complex in 3 mL of culture medium was then added to each well. Media was replaced after 48 hours with complete EGM-2. Transfection efficiency was verified by western blotting: cells were lysed in sample buffer containing DTT and LDS, boiled, and loaded onto a gel and separated with electrophoresis. Proteins were transferred to a 0.45-µm PVDF membrane. Blots were quantified by measuring band intensity using ImageJ and normalizing relative expression to control conditions. Membranes were incubated with anti-CD44(1:50) or anti-Beta actin(1:400) and visualized with HRP-conjugated secondary antibodies.

### Adducin-γ knockdown

DsiRNA directed against adducin-γ (ADD3)^30^ was used to knock down protein expression in HCMEC/d3 cells (IDBT). Cells were plated in a 6-well plate at 250k per well. When cells reached 60-70% confluency, DsiRNA was added as follows. Protocols were adapted from IDBT. Briefly, DsiRNA and negative control DsiRNA were resuspended in nuclease free water at 100 µM. The stock solutions were diluted to 5 µM working solution consisting of 5x siRNA buffer and nuclease free water. The working solution was diluted with transfection media to a 250nM concentration and incubated for 5 minutes at room temperature. Concurrently, Dharmafect was mixed with transfection medium and incubated for 5 minutes at room temperature. The solutions were mixed and incubated for another 5 minutes prior to adding the complex to complete EGM-2 to yield a final DsiRNA concentration of 25nM. 2 mL was added per well of cells and incubated for 24 hours before replacing with complete EGM-2. DsiRNA-mediated knockdown efficiency was verified using western blotting. Cells cultured in a well plate for five days following plating were lysed for the western blots, since this period matched the timing of permeability testing.

### RhoA activation and inactivation

In order to constitutively activate RhoA vessels at day 4 were perfused with Rho Activator II (Cytoskeleton) at a concentration of 2 µg/mL for four hours prior to use in permeability tests or fixation for immunocytochemistry. In order to assess the effects of RhoA inactivation, vessels were perfused with Rho Inhibitor I (Cytoskeleton) at a concentration of 2 µg/mL for four hours prior to use in permeability tests or immunocytochemistry.

### ELISA-based quantification of small GTPase activity

Commercial ELISA kits were purchased from Cytoskeleton (G-LISA) to quantify RhoA and Rac1 activation (Cytoskeleton). For both cases, cell lysates were prepared using Cytoskeleton’s protocols. Following exposure to flow or static conditions, the collagen/hyaluronan hydrogel was removed from the device and washed in ice cold 1x PBS for 30 seconds. 50 µL of ice-cold cell lysis buffer with 1x protease inhibitor was injected into vessels and collected in a microcentrifuge tube. The cell lysate solution was spun at 10,000 g for 1 minute at 4°C to pellet cell debris. The supernatant was collected and snap frozen in liquid nitrogen, reserving a small amount for protein quantification using Precision Red (Cytoskeleton). Prior to measuring GTPase activity, samples were thawed in a room temperature water bath and equilibrated to 1 mg/mL for RhoA and 0.5 mg/mL for Rac1 samples.

### Activated RhoA immunoprecipitation

Activated RhoA pulldown kits were purchased from Cytoskeleton. Sample lysates were prepared similar to the protocol used for G-LISAs. Samples were processed using Rhotekin-conjugated beads provided in the kit. 200 µg of protein was incubated with 15 µL of the beads, with 50 µg reserved for quantifying total RhoA present in the sample. Following pulldown, samples were denatured in Lamelli buffer and separated using electrophoresis. 10 µL of each sample was loaded per well on a 4-12% tris glycine gel in MOPS buffer. The gel was transferred onto a 0.2-µm PVDF membrane. Following transfer, the membrane was probed using the iBind flex kit and a primary anti-RhoA antibody (Cytoskeleton) at 1:200 and HRP-conjugated secondary (1:4000). Blots were quantified by measuring band intensity with ImageJ and normalizing relative expression to control conditions.

### ZO-1 immunoprecipitation

Immunoprecipitation of ZO-1 was performed using cell lysates extracted from cell monolayers cultured on the collagen/HA hydrogels exposed to 24hrs of fluid shear stress applied by a cone and plate rheometer. The gels were removed from the rheometer and washed immediately in ice cold PBS. The gels were then submerged in ice cold lysis buffer (CST) and sonicated. Gels were then spun down at 14,000 g for 10 minutes at 4°C. Supernatants were removed and snap frozen reserving a small aliquot for protein quantification. Samples were equilibrated at 0.5 mg/mL protein concentration using ice cold lysis buffer after thawing in a room temperature water bath. Samples were then loaded on a tris-acetate gel and separated using electrophoresis in tris-acetate buffer. Protein was transferred to a 0.45-µm PVDF membrane. The membrane was incubated overnight with ZO-1(CST) (1:2000), ADD3 (Abcam)(1:400), or Spectrin-αII (ABcam)(1:1000) antibodies and visualized with HRP-conjugated secondary antibodies (1:4000).

### Statistical Analysis

The open source statistics package, R, was used to perform all statistical calculations. Data sets were tested for normality with Shapiro-Wilk tests prior to testing for significance. One and two-way ANOVA tests followed by Tukey HSD post-hoc comparisons were used to evaluate significant differences between multiple conditions. Paired t-tests were used to compare fold change of ADD3/ZO1 and Spectrin/ZO1 with shear stress in the immunoprecipitation experiments. All other two sample comparisons were made using Student’s t-tests. Each statistical test used sample numbers greater than or equal to 3 unless otherwise noted and p < 0.05 was considered significant. All error bars indicate standard deviation of the mean.

## Results

### Fluid shear stress affects tight junctions and barrier integrity

In order to gain a more complete understanding of the effect of shear stress on barrier function, a range of shear stress levels was applied to the 3D BBB model for a period of four days. Four different levels of shear stress (static control, 0.18, 0.35, and 0.7 dyn/cm^2^) were exerted by altering the volumetric flow rate of culture medium perfusing through the 3D vessels, which are depicted in Figure 1A. These magnitudes were chosen to represent the lower end of the shear stress range present in the brain vasculature^31^, which previous measurements have shown is substantially heterogeneous^32, 33^. Each vessel was submerged in culture medium during these experiments to assure equal access to nutrients and oxygen regardless of perfusion rate. Three separate assays evaluated barrier integrity: immunocytochemistry, FITC-dextran permeability assays, and transendothelial electrical resistance (TEER) measurements. The former two assays were conducted at the end of the perfusion period, but TEER measurements could be performed daily without disrupting the vessels. Fig 1B-E show the effect of varying magnitudes of fluid shear stress on the morphology of the endothelial cells lining the *in vitro* vessel and localization of ZO-1, a tight-junction associated scaffolding protein^34^. After 4 days in culture, static vessels (Fig. 1B) exhibited a substantial amount of perinuclear ZO-1 staining, with irregular localization to the junctions. Vessels exposed to low levels of shear stress (0.18 and 0.35 dyne/cm^2^) had a similar response, as evidenced in Figures 1C and 1D, respectively. In contrast, the vessel exposed to 0.7 dyn/cm^2^, indicated clear localization of ZO-1 to the cell-cell junctions without any gaps in the endothelial monolayer (Fig. 1E).

**Figure 1.**
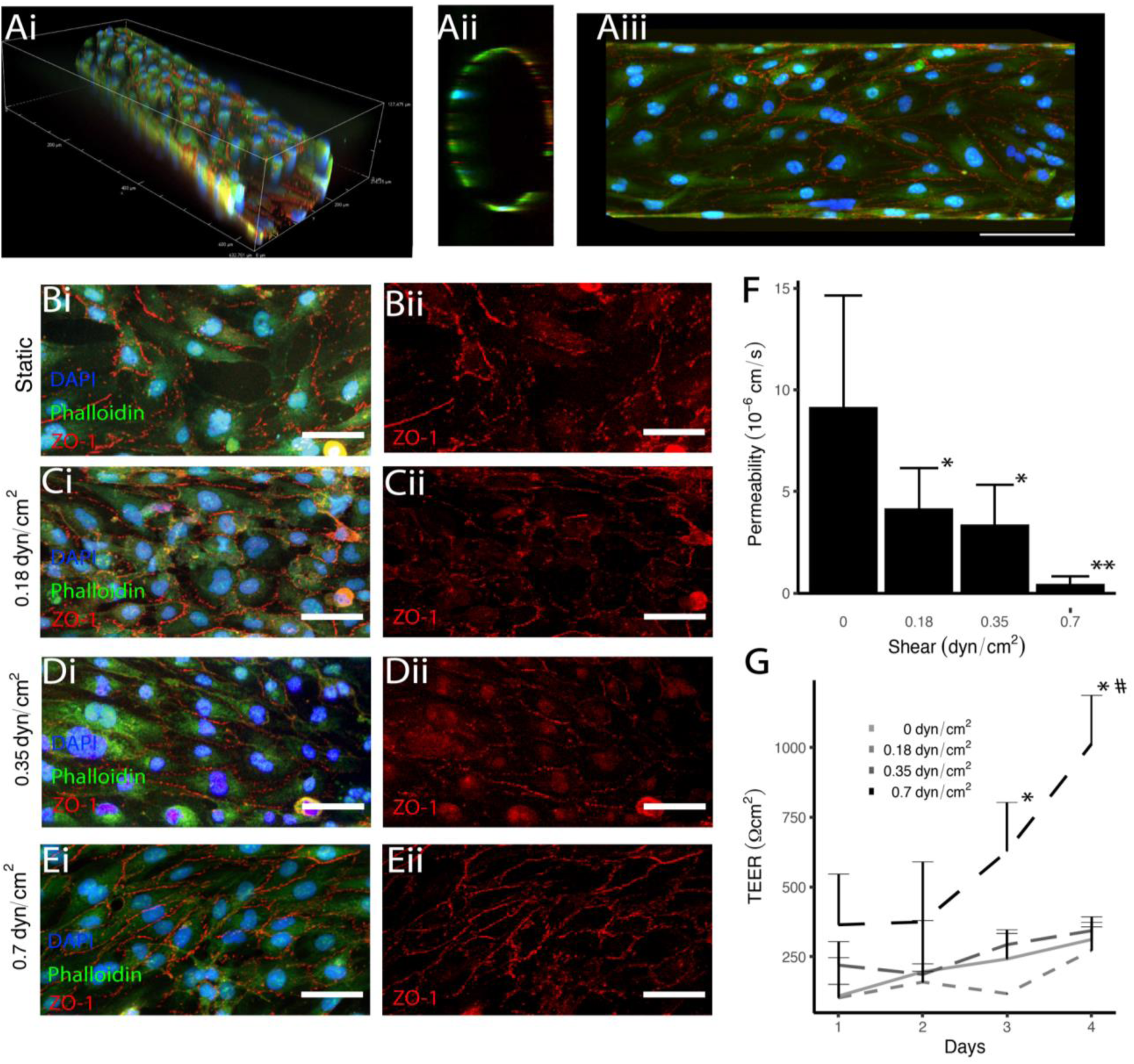
Fluid Shear stress affects tight junctions and barrier integrity. **A** Fluorescent images of HCMEC/D3-seeded channels cultured for 4 days in 3D (i), cross-section (ii), and maximum projection in the x and y plane(iii) scale=100μm. **B** Static, **C** 0.18 dyn/cm^2^, **D** 0.35 dyn/cm^2^, **E** 0.7dyn/cm^2^ images showing DAPI (blue), phalloidin (green), and anti-ZO-1 (red) (isolated in ii) staining of the vessel wall. scale=50 μm **F** Permeability coefficient of channels exposed to static or shear conditions for 4 days measured with 4kDa dextran, *denotes p < 0.05 and ** denotes p < 0.001 compared to static condition. **F** TEER measurements for channels exposed to static or shear conditions for 4 days, *denotes p < 0.05 compared to Day 1 value for each condition. # indicates p<0.05 compared to Day 4 values for each condition

The permeability measurements quantitatively validated the observations yielded by immunocytochemistry. Figure 1F indicates that only the vessels exposed to 0.7 dyn/cm^2^ shear stress resulted in a significantly lower permeability compared to lower shear stress magnitudes and the static control. Similarly, the TEER measurements found a significant increase in barrier integrity only in vessels exposed to 0.7 dyn/cm^2^ at days 3 and 4 following seeding of the cells within the hydrogel (Fig. 1G). TEER was also used to evaluate whether the presence of astrocytes in the surrounding hydrogel were required for barrier formation. Supplemental Figure 1A-B indicates no significant difference in FITC-dextran permeability and TEER values in hydrogels containing astrocytes. These findings further emphasize the importance of fluid shear stress in barrier regulation in cerebral vasculature.

### Hyaluronan mediates formation of the endothelial barrier

Another means of interrogating the effect of shear stress on barrier function involves evaluating the dynamics of endothelial tight junctions after ceasing perfusion. These experiments also have clinical relevance due to the occlusion of cerebral blood flow during ischemic stroke. Vessels were exposed to 0.7 dyn/cm^2^ for four days prior to stopping flow (at time = 0) and evaluating permeability at 20, 60, 120, and 240 minutes. As Figure 2A indicates, the permeability of the 3D vessels significantly increased at the 120 and 240-minute timepoints following removal from flow. These experiments were also conducted in the absence of hyaluronan (HA), to determine the importance of matrix formulation on flow-mediated barrier integrity. As Figures 2A and 2B show, vessels fabricated in collagen-only hydrogels exhibited a significantly higher permeability at the time when flow stopped, and there was no significant change in permeability in the four hours that followed. These results suggested that the presence of HA contributed to flow-mediated barrier formation and maintenance.

**Figure 2.**
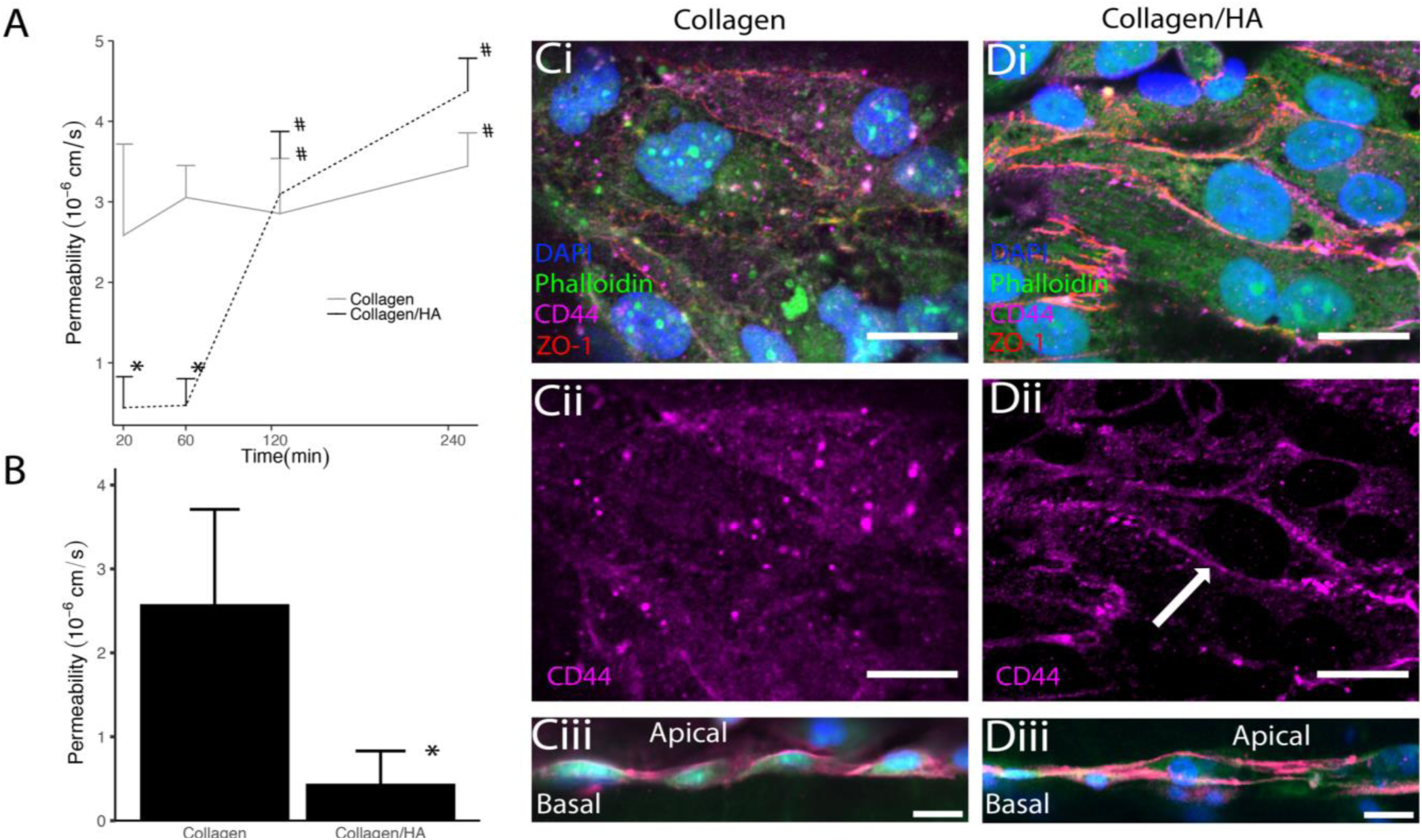
ECM composition affects barrier integrity and CD44 localization. **A** Permeability measurements taken after 4 days of perfusion and for 4 hours after flow stoppage. **B** Permeability coefficients measured using 4kDa dextran after 4 days of perfusion for channels in a collagen or collagen/HA hydrogel. **C** Localization of CD44 in vessels patterned in a collagen only hydrogel and stained for DAPI (blue), phalloidin(green), anti-ZO-1 (red), and anti-CD44 (magenta) (isolated in ii) in the vessel wall (i and ii) and in cross-section (iii). Scale: 20μm. **D** Localization of CD44 in vessels patterned in a collagen/HA only hydrogel. Scale: 20μm. * indicates p<0.05 between collagen and collagen/HA scaffolds at each time point, and # indicates p < 0.05 compared to the 20-minute time-point.

Because CD44 is one of the main transmembrane receptors for HA, experiments were conducted to determine whether the receptor was involved in mechanotransduction of fluid shear stress in the cerebral endothelial cells. First, immunocytochemistry was performed on vessels patterned in both collagen/HA and collagen only scaffolds and exposed to four days of 0.7 dyn/cm^2^ shear stress. Figures 2C-D provide images of these vessels, and show that CD44 localizes to the cell-cell junctions in vessels patterned in collagen/HA scaffolds, whereas the receptor is more diffuse throughout the cytoplasm and does not localize strongly to cell-cell junctions in the collagen only hydrogels. Side views of the vessels in collagen only hydrogels reveal CD44 localized to the apical sides of the endothelium where there is likely a robust glycocalyx layer (Fig. 2Ciii), though positive staining is observed in both apical and basal sides in collagen/HA hydrogels, suggesting that binding of the receptor to the extracellular matrix is crucial for the observed flow response (Fig 2Diii).

### CD44 mediates barrier formation in response to shear stress

In order to further interrogate the role of CD44 in flow mechanotransduction, HCMEC/D3 cells were genetically modified using the CRISPR/Cas9 system to both knockout and upregulate the expression of the receptor. A CRISPR/Cas9 plasmid containing a scrambled guide sequence was used as a control for these experiments. Double nickase plasmids were used to knockout CD44^35^, and the SAM activation system was used to constitutively increase expression^36^. Western blotting validated the effect of gene editing on CD44 expression (Figs. 3A-B). Having measured CD44 expression levels, four separate conditions were evaluated using immunocytochemistry, FITC-dextran permeability tests, and TEER measurements: (i) scrambled control cells in collagen/HA hydrogels exposed to flow, (ii) CD44-upregulated cells in collagen only hydrogels exposed to flow, (iii) CD44-upregulated in collagen/HA hydrogels in static culture, and (iv) CD44 knockout cells in collagen/HA hydrogels exposed to flow. The purpose of these conditions was to determine whether CD44 was necessary for flow-mediated barrier formation and whether increasing the expression of CD44 could recover barrier function in the absence of HA or flow. Immunocytochemistry provided insight into the morphology and expression of tight junctions; Figures 3C-F provide composites of DAPI and GFP with ZO-1 (i) as well as isolated ZO-1 images (ii). Vessels treated with scrambled guide sequences were not affected by transfection, ZO-1 is observed at the cell-cell junctions with minimal perinuclear localization (Fig. 3C). The transformed cells did not exhibit different barrier integrity compared to normal cells determined by permeability and TEER measurements (Supplementary Figure 2). Upregulation of CD44 in static collagen/HA hydrogels was not able to recover barrier formation observed in the presence of flow, as evidenced by a disrupted monolayer and diffuse ZO-1 staining (Fig. 3D). Similarly, constitutive activation of CD44 in collagen scaffolds exposed to flow exhibited limited ZO-1 localization to the cell-cell junctions (Fig. 3E), demonstrating that expression of CD44 in the absence of HA could not recover function. Finally, CD44 knockout cells in collagen/HA scaffolds exposed to flow resulted in discontinuous localization of ZO-1 to the cell-cell junctions and gaps within the monolayer (Fig. 3F). FITC-dextran permeability assays (Fig. 3G) and daily TEER measurements (Fig. 3H) confirm these immunocytochemistry observations. In both assays, the scrambled control had significantly higher barrier function compared to the other conditions. Overall, these results suggest that all three elements: ZO-1 expression, HA in the extracellular matrix, and shear stress are required for flow-mediated barrier formation.

**Figure 3:**
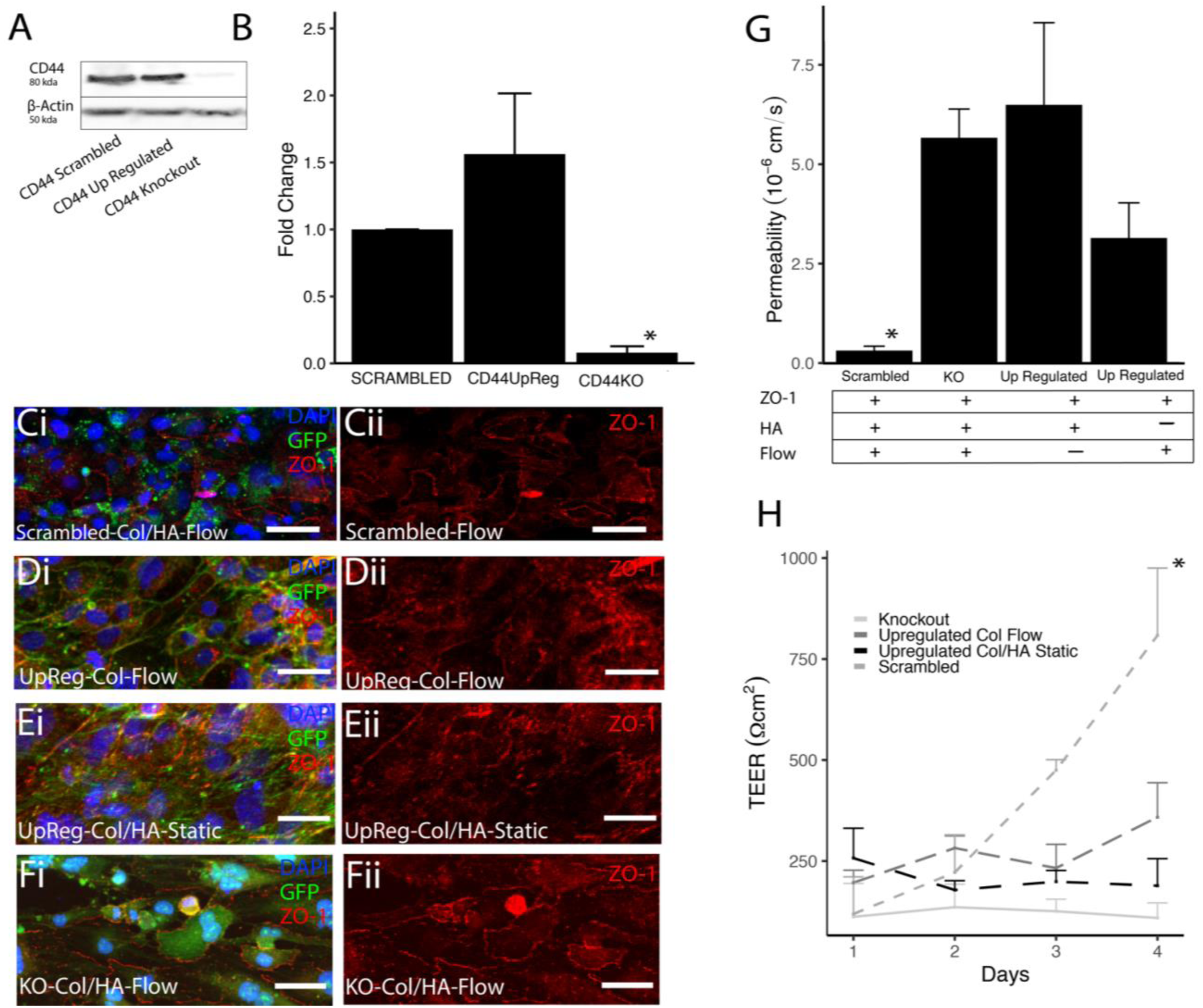
Expression of CD44 regulates barrier formation. **A** Western blot of CD44 and beta-actin in CRISPR-modified cells. **B** Ratio of intensities of CD44 and beta-actin signals normalized to scrambled levels. * indicates p<0.05 compared to scrambled and upregulated conditions **C-F** Fluorescent images of channels stained with DAPI (blue), GFP (green), and anti-ZO-1 (red) (isolated in ii) for four conditions: scrambled control cells in collagen/HA hydrogels exposed to flow (**C**), upregulated cells in collagen hydrogels exposed to flow (**D**), upregulated cells in collagen/HA hydrogels in static conditions (**E**), and knockout cells in collagen/HA hydrogels exposed to flow (**F**). **G** Permeability coefficients of the channels seeded with transfected cells after 4 days of culture * denotes p <0.05 compared to all conditions. **H** TEER measurements taken over the course of 4 days for different conditions, * indicates p < 0.05 compared to Day 1 timepoint for all conditions. Scale: 50μm

### Small GTPases regulate barrier integrity

Given that previous studies have found CD44 activation affects small GTPases associated with cell mechanotransduction^37^, studies were conducted to determine the effect of shear stress on RhoA and Rac1 activation within the 3D BBB model. Although the effect of RhoA activation on the blood-brain barrier has been extensively studied^8, 16, 21, 38–41^, its response to fluid shear stress in cerebral endothelial cells has yet to be studied. Here, vessels were treated with a Rho inhibitor prior for four hours prior to stopping flow or treated with a Rho activator in the last four hours of the four-day perfusion. Immunocytochemistry shows that modulating RhoA activation has a substantial effect on the morphology of the endothelial cells. Figure 4A shows strong ZO-1 localization to the junction following 240 minutes of no flow. Additionally, the membrane protein adducin**-γ** was also localized to the junction in this condition (Fig. 4A, iii). In contrast, exposure to the Rho activator resulted in barrier disruption and more perinuclear staining of the ZO-1 (Fig. 4B). FITC-dextran permeability testing with vessels perfused with the Rho inhibitor exhibited no significant increase in permeability, as shown in Figure 4C. Additionally, the permeability at the end of the four-day perfusion was significantly higher in vessels treated with the Rho activator (Fig. 4D).

**Figure 4:**
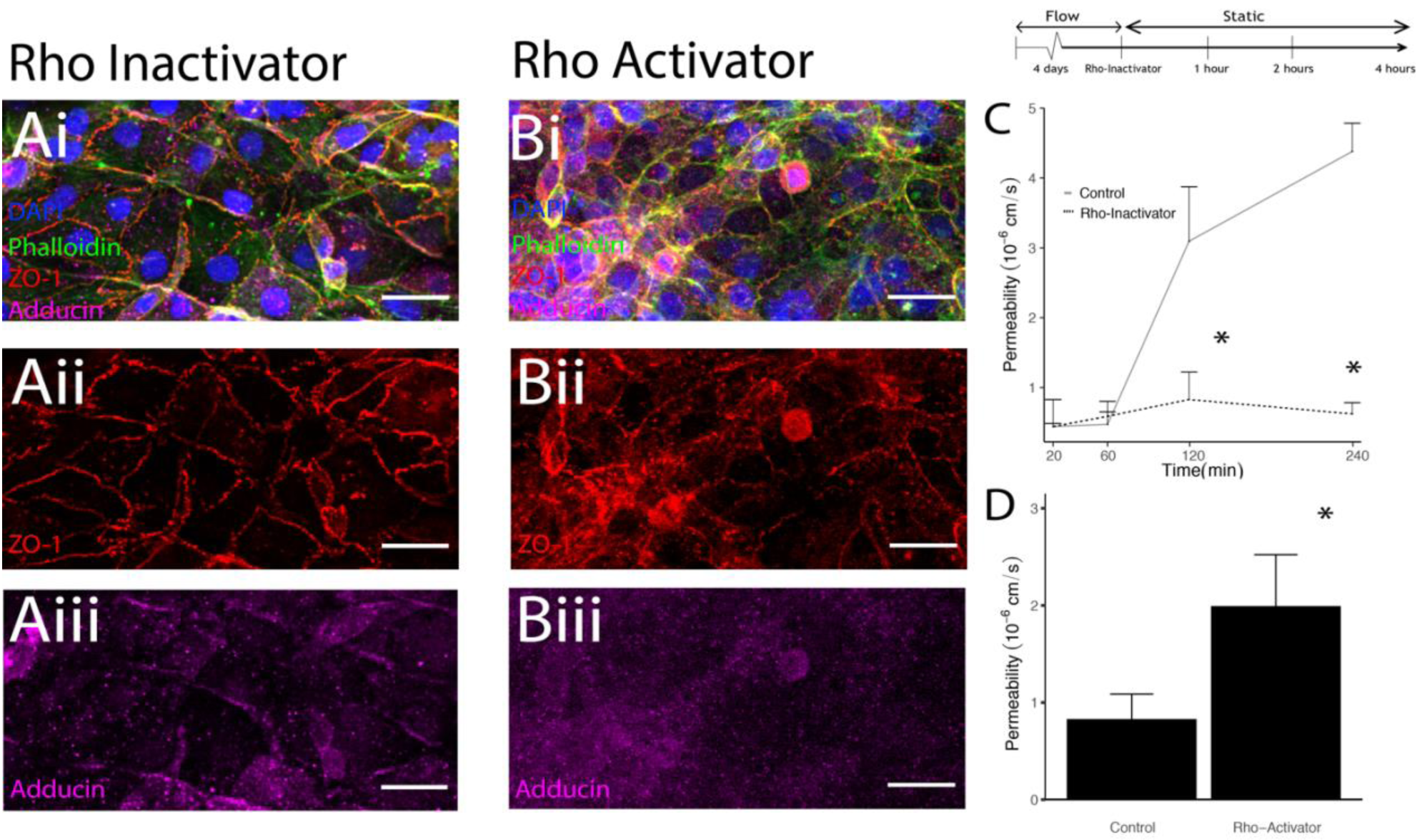
Fluid shear stress mediates Rac1 and RhoA activation. **A-B** Fluorescent images of vessels stained with DAPI (blue), ZO-1(red) (isolated in ii), and adducin (magenta) (isolated in iii) for Rho Inactivator (**A**) and Rho Activator (**B**). Scale: 50μm **C** Permeability coefficients of channels perfused for 4 days then incubated with a Rho inactivator for 4 hours prior to flow stoppage. * indicates p<0.05 compared to the control condition for each time point. **D** Permeability coefficients of channels perfused for 4 days then exposed to a Rho Activator for 4 hours prior to measurement. * indicates p < 0.05.

ELISAs and immunoprecipitation assays provided further insight into the effect of flow on the activation of small GTPases. Due to the amount of protein required for these assays, the 3D BBB model was altered for these experiments. A larger diameter vessel (1-mm), which has been previously used for high Reynolds number studies^26^, was endotheliaized and exposed to 2.1 dyn/cm^2^ of shear stress. Figure 5 shows that application of 2.1 dyn/cm^2^ significantly reduced the level of RhoA activity, as measured by the Rhotekin-pulldown (Fig. 5A) and ELISA (Fig. 5B) assays. Both assays indicated a reduction of nearly 50% compared to static controls. An ELISA was also conducted for Rac1, and found that Rac1 activation was significantly increased at the 6-hr and 24-hr time points following the start of perfusion. These assays indicate that flow results in differential expression of RhoA and Rac1: shear stress significantly decreases RhoA activation while significantly increasing Rac1 activation.

**Figure 5.**
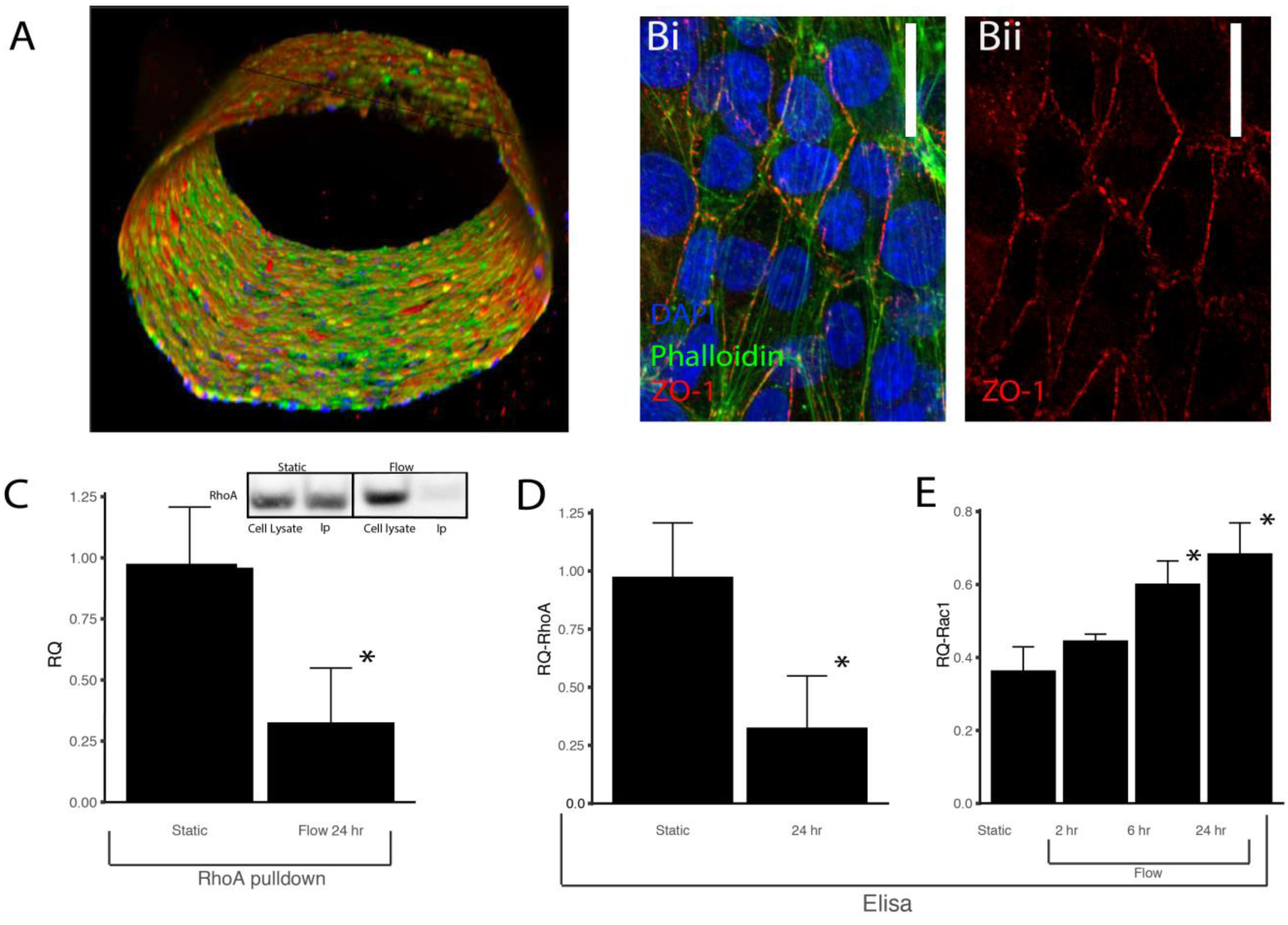
Fluid Shear Stress mediates Rac1 and RhoA activation. **A** 3D confocal scan of the 1-mm diameter vessel. **B** HCMEC/D3 stained for DAPI (blue), phalloidin (green), and anti-ZO-1 (red) (isolated in ii) after exposure to 2.1 dyne/cm^2^ in the enlarged vessel. **C** RQ of activated RhoA to total RhoA isolated using immunoprecipitation as measured by western blot. **D** Relative intensity (RQ) of RhoA activation in channels exposed to flow measured with ELISA. **E** Relative intensity (RQ) of Rac1 activation in channels exposed to flow measured with ELISA. *indicates p<0.05 compared to the static condition.

### Adducin-γ localization is a downstream effect of Rac1 activation

Given that adducin-γ localized to the junctions in vessels exposed to the Rho inhibitor, experiments were conducted to determine the importance of this membrane protein to flow-mediated barrier formation. In order to determine whether adducin-γ expression was necessary for barrier integrity, siRNA was used to knockdown levels of adducin-γ in the HCMEC/D3 cells (Figs. 6A-B). Knockdown efficiency was measured 5 days after transfection to ensure that *in vitro* vessels seeded with treated endothelial cells would express reduced levels of adducin**-γ** during the perfusion period. Permeability testing of vessels containing the knockdown cells exhibited significantly higher permeability compared to scrambled controls (Fig. 6C). As Figures 6D-E indicate, the localization of adducin-γ to the junctions was significantly different in vessels exposed to 0.7 dyn/cm^2^ shear stress compared to static controls. In order to determine whether a tight junction-associated complex incorporating adducin-γ formed in response to flow, ZO-1 immunoprecipitations were performed. Similar to the ELISAs used to evaluate small GTPase activity, a substantial amount of protein was also required for these experiments. Therefore, HCMEC/D3 cells were seeded on the surface of collagen/HA hydrogels and placed in a cone and plate rheometer to apply a steady shear stress of 0.7 or 2.1 dyn/cm^2^. Immunocytochemistry verified that the monolayer exhibited a difference in ZO-1 localization in response to shear stress (Supplementary Figure 3), indicating that cells seeded on flat gels responded similarly to cells seeded within a 3D vessel. As Figures 6F-G show, the application of shear stress resulted in significantly higher levels of adducin-γ in the ZO-1 pulldown compared to static controls. Significantly increased levels of spectrin-αII bound to the ZO-1 complex were also measured in response to shear (Supplementary Figure 4), consistent with previous findings that spectrin binds to adducin^42^. Overall, these results indicate that adducin-γ localization is an integral downstream result of flow-mediated CD44 activation and differential activation of Rac1 and RhoA.

**Figure 6:**
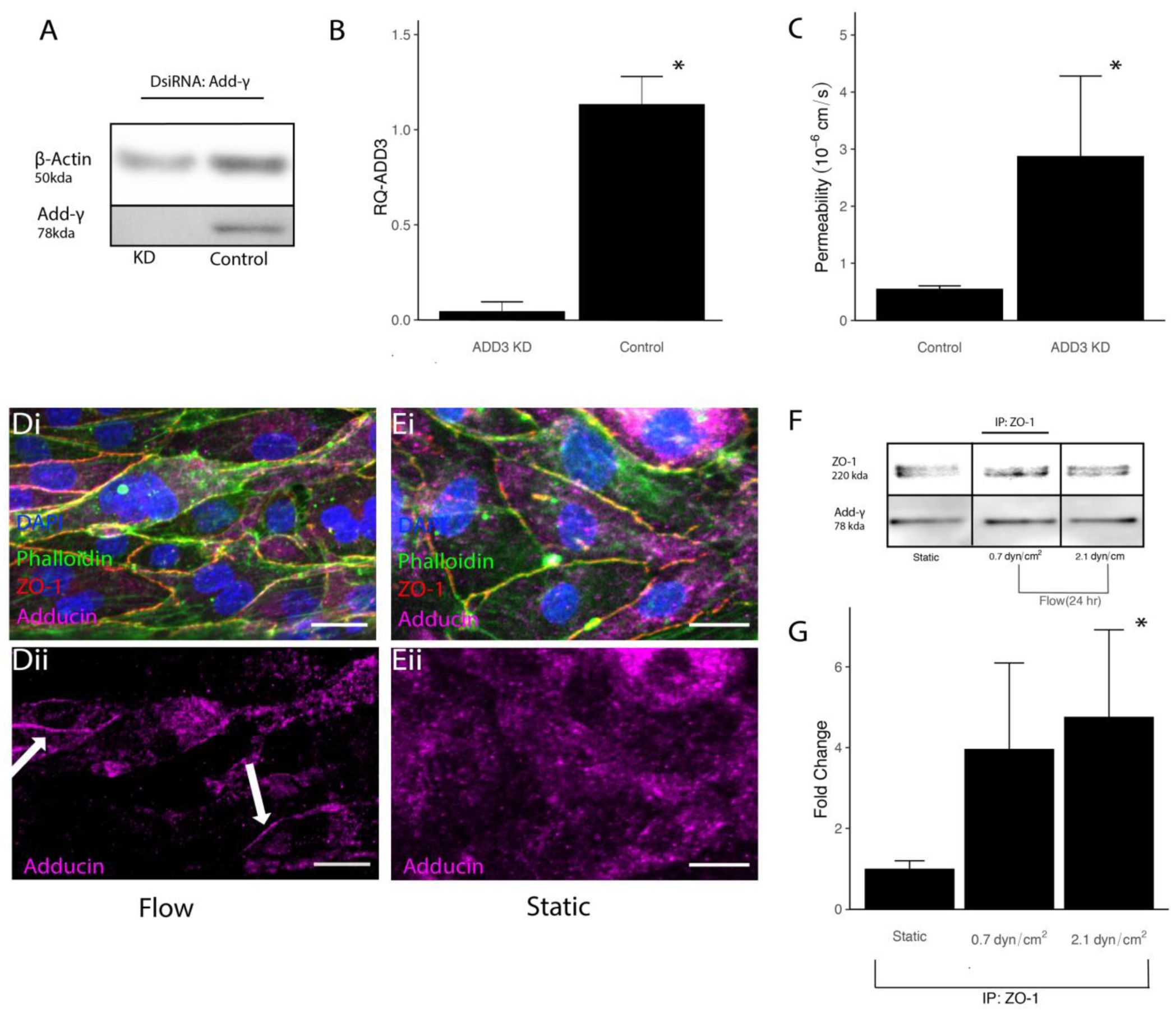
Adducin-γ regulates tight junctional stability through association with ZO-1. **A** Western blot of cells treated with DsiRNA targeting ADD3. **B** Relative intensity of ADD3 in DsiRNA and control cells normalized to beta-actin. **C** Permeability coefficients measured in channels seeded with ADD3 Knockdown (KD) HCMEC/d3 cells* indicates p<0.05 compared to control. **D-E** Fluorescent images of vessels stained with DAPI (blue), phalloidin (green), ZO-1 (red), and ADD3 (magenta) (isolated in ii) for flow (**D)** and static (**E**) conditions. **F** Immunoprecipitation of the ZO-1 complex from cells exposed to static and flow conditions. **G** Relative intensity of ADD3 bound to ZO-1 isolated using IP and measured with western blot. Scale: 25μm, * indicates p<0.05 compared to static

## Discussion

Our results identify a previously unreported signaling transduction mechanism that mediates blood-brain barrier formation in response to fluid shear stress. Figure 7 summarizes the components of this pathway. CD44 acts a mechanosensory and localizes to junctions in response to shear stress, activating Rac1 while inhibiting RhoA, which induces a complex including ZO-1, adducin-γ, and spectrin-αII at the cell-cell junctions. Our findings suggest that a requisite level of shear stress is needed to activate this pathway, given that levels of shear stress less than 0.7 dyn/cm^2^ were unable to significantly increase barrier formation and maintenance during perfusion. Moreover, CRISPR-mediated alteration to CD44 levels suggest that all three components: hyaluronan, CD44, and shear stress all bolster barrier integrity in the in vitro 3D BBB model used. Our study concludes that this CD44-mediated mechanism is integral to blood-brain barrier mechanotransduction, identifying an alternative mechanosensors of shear stress^14^.

**Figure 7:**
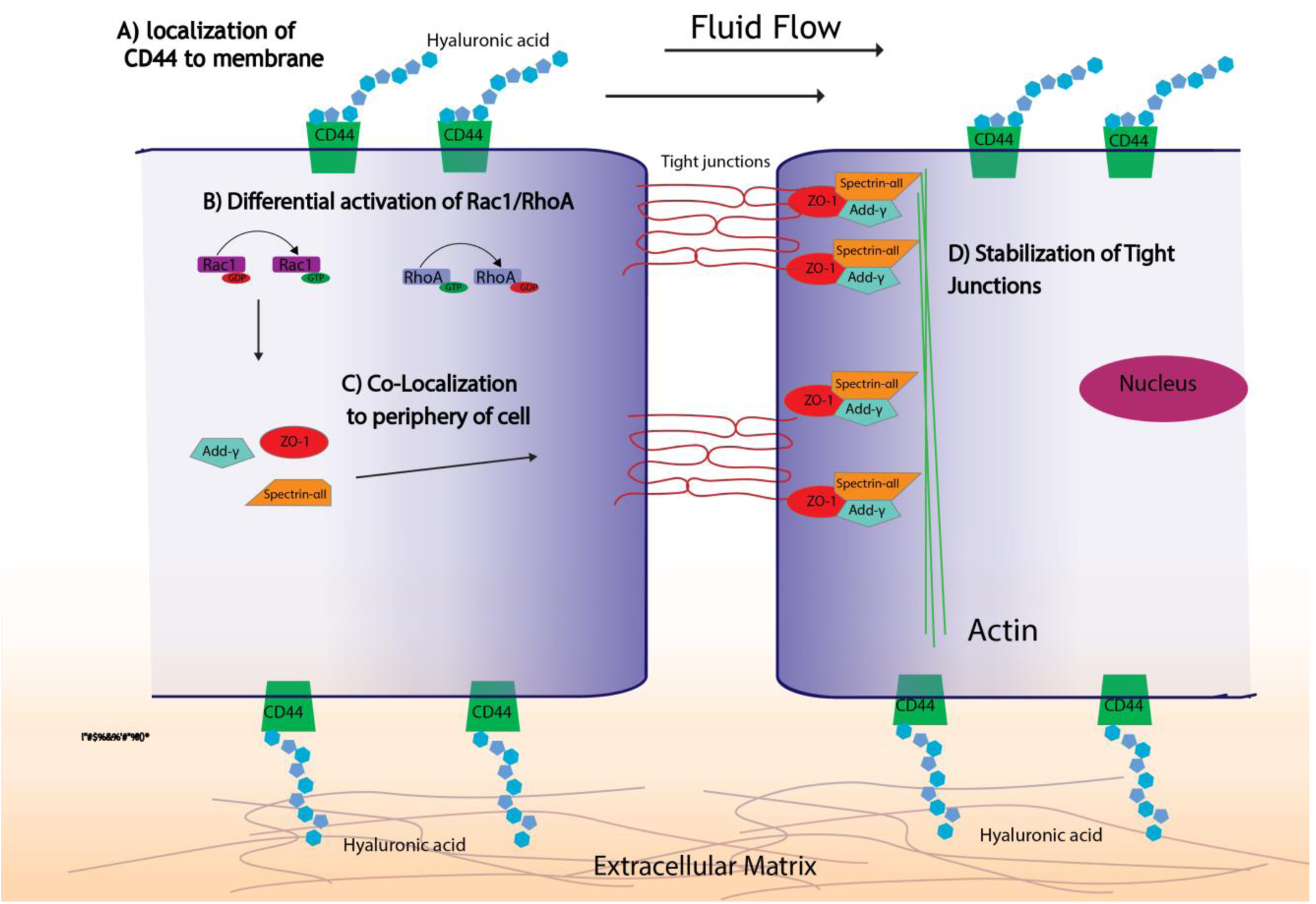
Mechanosensitive CD44 helps stabilize tight junctions through activation of Rac1. **A** CD44 bound to HA at the surface of endothelial cells transduces fluid flow. **B** The application of shear stress activates Rac1 while inactivating RhoA. **C** Activation of Rac1 leads to the recruitment of ADD3 and ZO-1 to the cell-cell junction. **D** ADD-3 and ZO-1 form a binding complex with spectrin-αII that stabilizes tight junctions that give rise to the blood-brain barrier.

The role of CD44 in mechanotransduction has been well-studied^43^, but its importance for sensing shear stress has yet to be investigated in detail. A previous study using bovine aortic endothelial cells found that inhibiting hyaluronan synthesis negated flow-mediated cell spreading on soft (100 Pa) two-dimensional substrates^44^, suggesting that CD44 may mediate acto-myosin contractility in the presence of shear stress. Here, the importance of CD44 was initially revealed due to the difference between vessels fabricated in collagen-only and collagen/HA hydrogels. There was a significantly lower permeability in collagen/HA hydrogels where CD44 is present on both apical and basal sides, compared to collagen-only hydrogels where CD44 is only present on the apical side, likely due to the presence of the glycocalyx. Previous studies have demonstrated the importance of the glycocalyx in flow mechanotransduction, and vessels fabricated in collagen-only hydrogels still have a significantly lower permeability than static controls after perfusion. Yet, our results demonstrate that the presence of HA in the ECM yields a significantly lower permeability, suggesting that cell-matrix adhesions are important for mediating the flow response.

ELISA and pulldown studies indicated that shear stress also significantly increased the activation of Rac1 while attenuating the activation of RhoA, demonstrating that the activity of these small GTPases is affected by shear stress. The activation of RhoA has been shown to induce cell contractility and cytoskeleton restructuring, resulting in enhanced cell motility and disrupted barrier integrity^20, 21, 45^. Conversely, Rac1 has been identified as a key mediator of vascular integrity in systemic vasculature^12, 15^. Previous studies have shown that CD44 can activate these small GTPases^37, 46^, but the results presented here find that this interaction is important in mechanosensing. It is possible that CD44 may differentially regulate RhoA and Rac1 depending upon both mechanical and biochemical factors. A recent study found that the presence of small fragments of HA (< 4–8 kDa) resulted in CD44-mediated barrier breakdown^47^, suggesting that the molecular weight of HA may modulate the CD44 response. Overall, these results suggest that the effect of CD44 on downstream small GTPases can be altered by multiple factors.

Another crucial component of the signaling mechanism involves adducin-γ localization to tight junctions in the presence of shear stress. Adducin-γ has been shown in previous work to stabilize cell-cell junctions^48^ and is a ROCK substrate^42, 49^. Studies have found that adducin-γ is an important component of both adherens and tight junctions within endothelial cells, and that it mediates cAMP signaling^50^. Additionally, the increase in spectrin-αII bound to the tight junction complex in response to flow indicates a potential role in stabilization of the tight-junction complex. Previously studies have identified the ankyrin-adducin-spectrin complex, which binds to transmembrane proteins bridging the gap to the cytoskeleton^51^. Specifically, spectrin-αII is associated with the brain and its cleavage products are a potential biomarker for neurodegenerative disorders^52^. These findings indicate that that the association of adducin-γ and spectrin-αII may contribute to barrier function, yet additional studies are required to determine how the activity of RhoA and Rac1 affects the localization of these proteins in cell-cell junctions. Nonetheless, the knockdown studies suggest that adducin-γ is required to maintain blood-brain barrier integrity, since reducing its expression resulted in increased permeability and barrier disruption. Clinical studies have identified adducin-γ as a factor that can contribute to altered cerebral vascular function^53^, providing further evidence of its importance in maintenance of the blood-brain barrier.

Taken together, these results demonstrate a new mechanism by which fluid shear stress is transduced by endothelial cells within the blood-brain barrier. These findings point to new directions for therapeutic treatments to attenuate blood-brain barrier leakage during the “no-reflow” period following cerebral ischemia-reperfusion injury when attenuation of cerebral blood flow correlates with increased barrier breakdown and other pathologies associated with vascular degradation in the central nervous system^3^. Future work is required to investigate crosstalk between this pathway and previously studied mechanosensitive mediators of vascular integrity^6, 11, 12, 54^, and to determine ways in which this mechanism can be manipulated in vivo. Overall, the proposed mechanism not only provides targets for new therapeutic approaches, but also contributes to a mechanistic understanding of blood-brain barrier formation, specifically in response to fluid shear stress.

## Acknowledgments

This work was supported by an American Heart Asssociation Scientist Development Grant awarded to P.A.G. (17SDG33460432), National Institute of Health grant awarded to F.F (RO1AG057842). and National Science Foundation GRFP fellowship awarded to B.J.D.

## Supplemental Figures

**Supplemental Figure 1:**
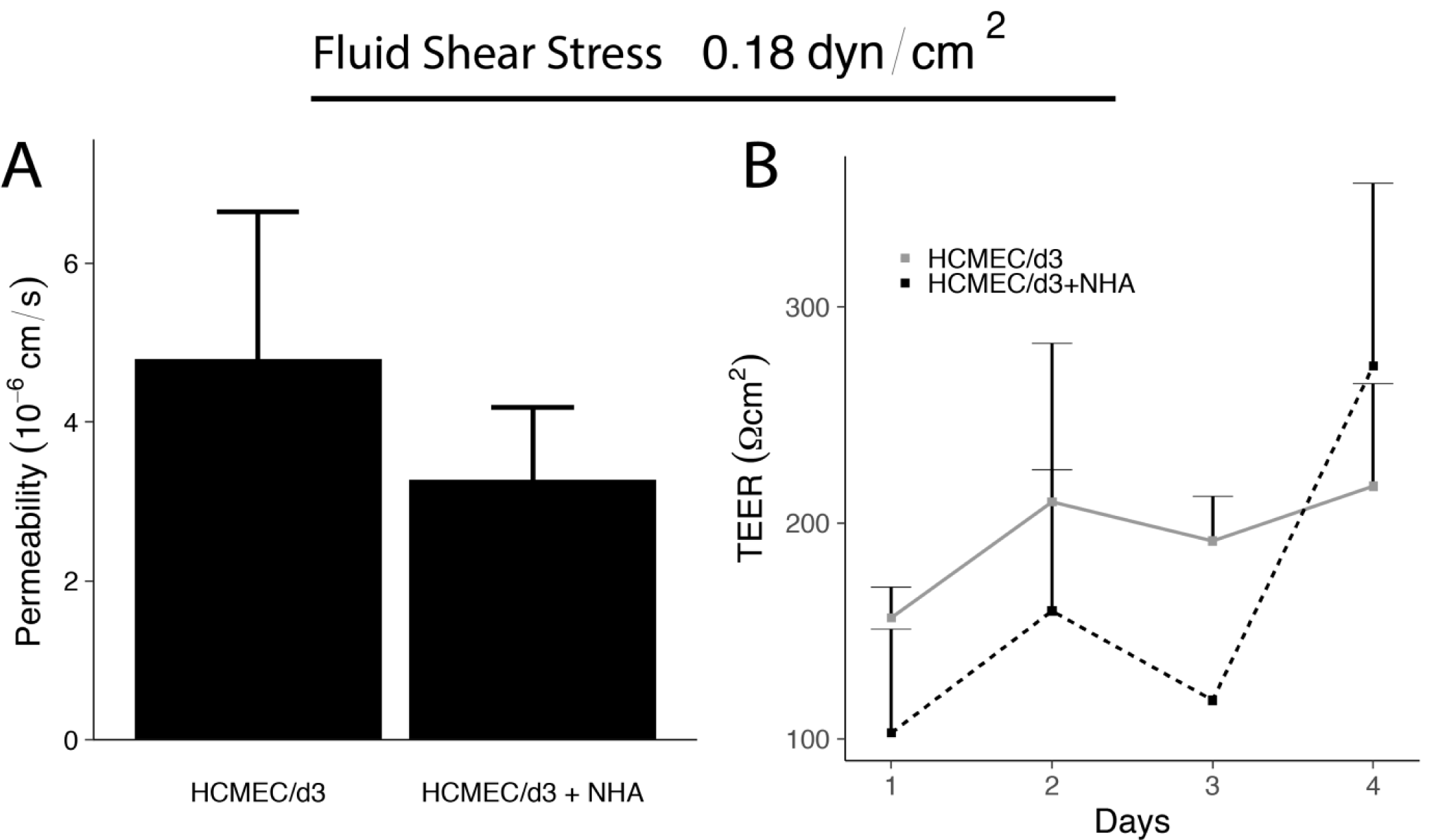
Contribution of astrocytes to barrier formation. **A** Permeability coefficient measured in channels seeded with HCMEC/D3 cultured with and without NHA after 4 days of perfusion. **B** TEER measurements in channels seeded with HCMEC/D3 cultured with and without NHA.

**Supplemental Figure 2:**
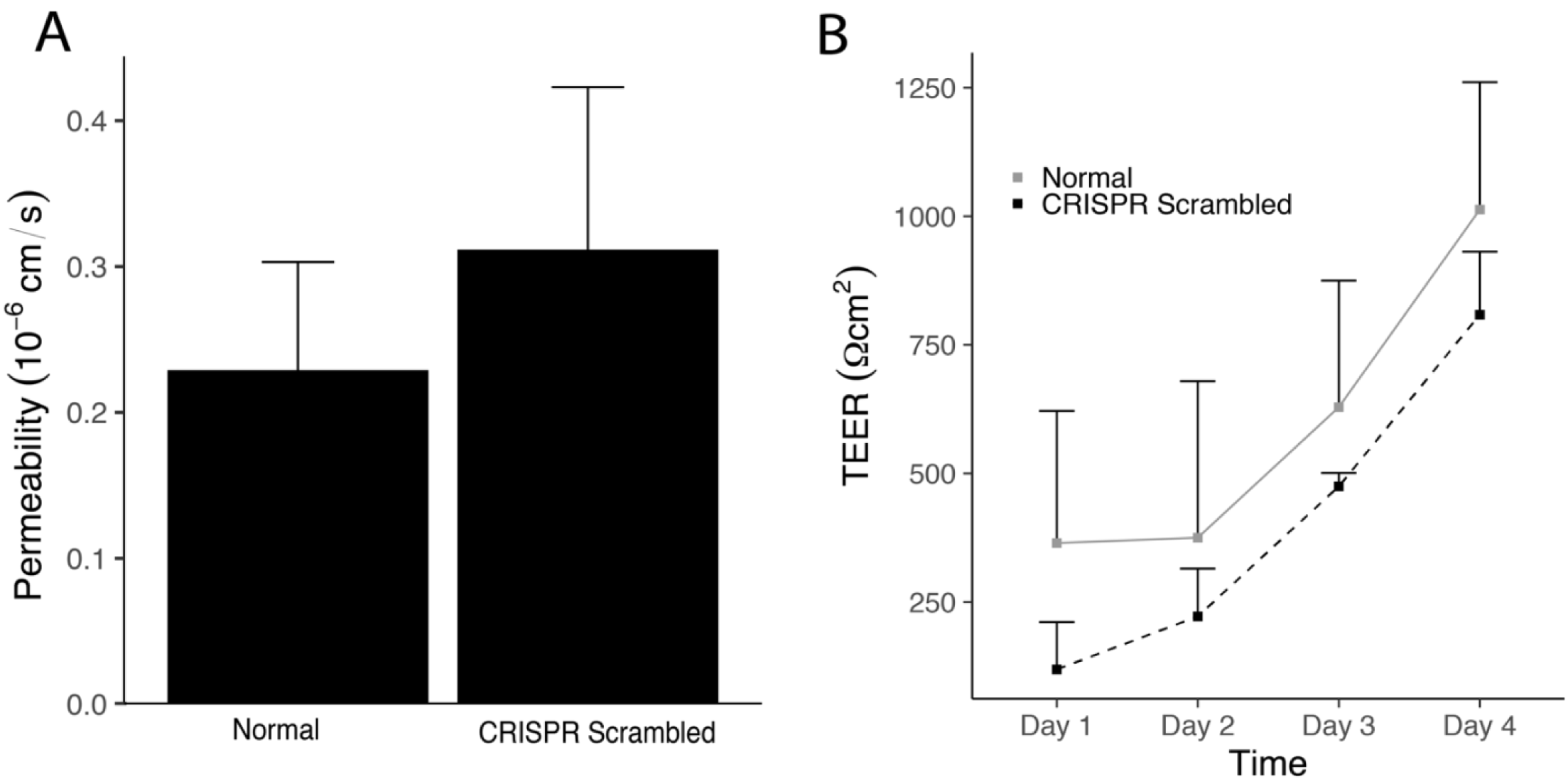
Barrier integrity of normal and transfected HCMEC/D3. **A** Permeability coefficients measured in channels seeded with HCMEC/D3 and exposed to 0.7 dyn/cm^2^ for four days comparing cells transfected with scrambled CRISPR/Cas9 plasmids with normal cells. **B** TEER measurements for channels seeded with both normal and transfected cells.

**Supplemental Figure 3:**
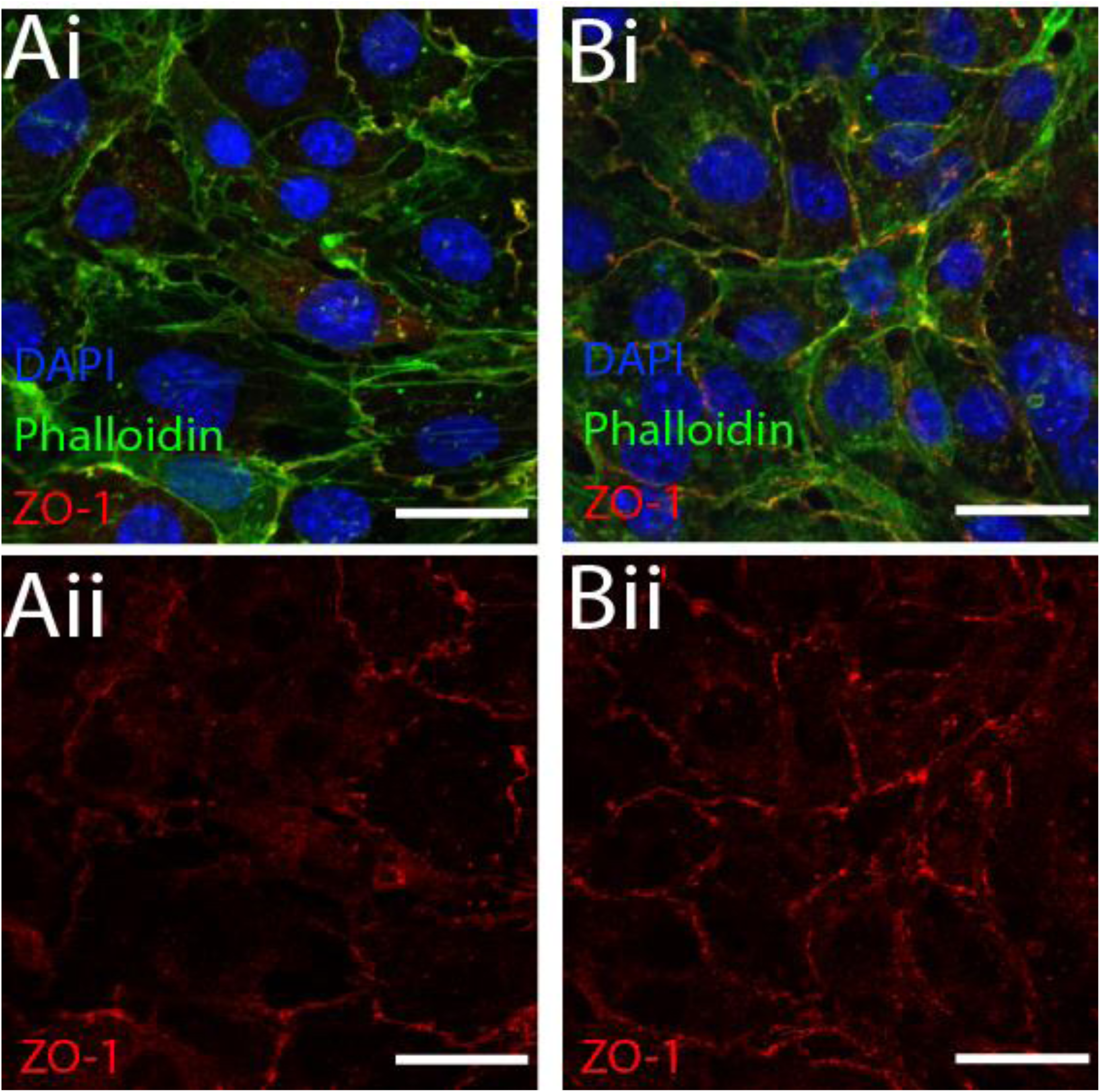
ZO-1 localization in cell monolayers used for immunoprecipitation experiments. **A-B** Monolayers stained for DAPI(blue), phalloidin(green), and anti-ZO-1(red) (isolated in ii) for static (**A**) and shear (**B**) conditions after 24 hours. Scale = 25 μm

**Supplemental Figure 4:**
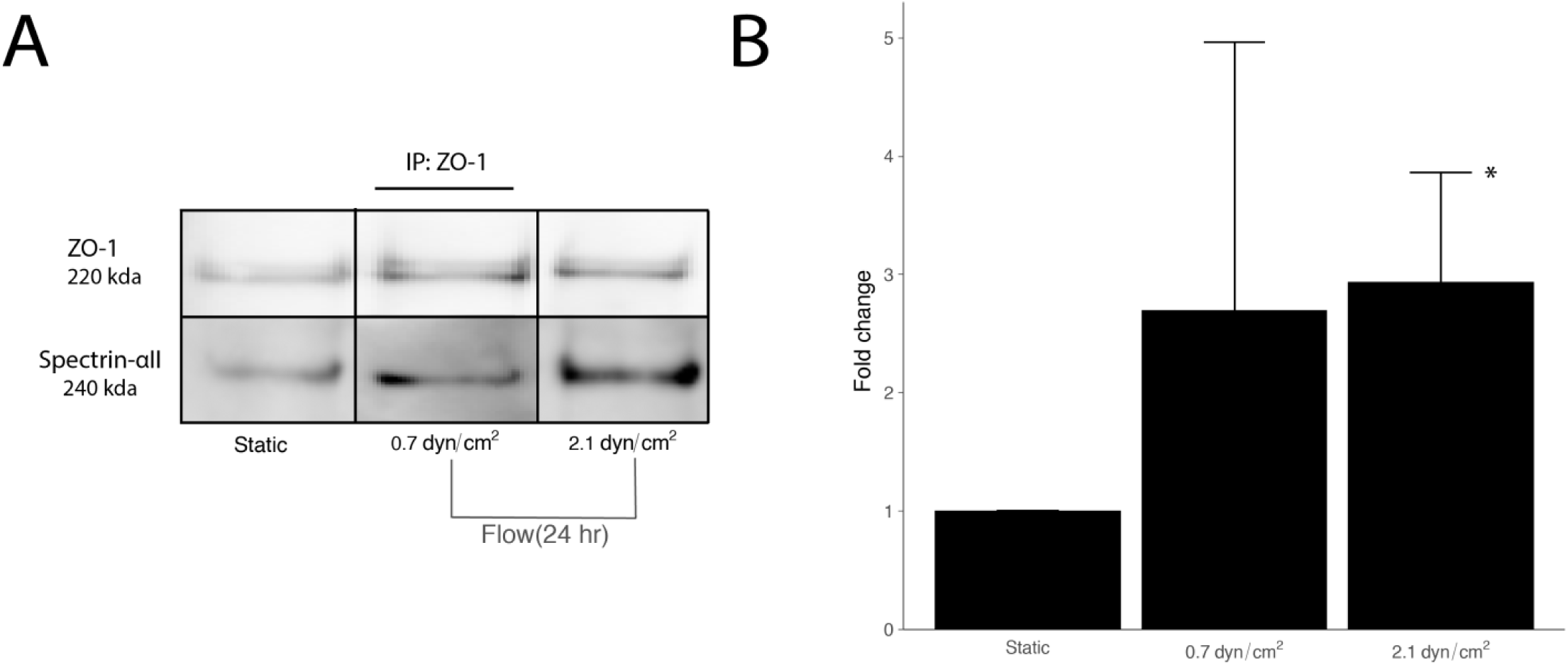
Probing ZO-1 pulldown for spectrin-αII. **A** Spectrin-αII isolated using IP from HCMEC/d3 seeded on monolayers exposed to 24 hrs of static or exposure to 0.7 or 2.1 dyn/cm^2^. **B** Fold change of the ratio of Spectrin-αII /ZO-1 normalized to static conditions. * denotes p < 0.05.

